# Protective Effects of Boric Acid Against LPS-Induced Inflammation and Apoptosis in a Primary Human Chondrocyte Model of Osteoarthritis

**DOI:** 10.64898/2026.08.13.744490

**Authors:** Mohammad Amin Yousefzadeh, Mahshid Azizi, Mohammad Hossein Nabian

## Abstract

Osteoarthritis is characterized by inflammation, chondrocyte dysfunction, and progressive cartilage degradation. Boric acid (BA), a physiologically relevant boron compound, has shown anti-inflammatory properties, but its effects on human articular chondrocytes remain unclear. This study investigated whether BA could protect primary human chondrocytes against lipopolysaccharide-induced inflammatory injury. Cell survival, membrane damage, apoptosis, inflammatory mediator production, and expression of genes related to inflammation and extracellular matrix degradation were assessed. BA improved chondrocyte survival and reduced membrane damage and apoptosis following inflammatory stimulation. It also suppressed inflammatory and matrix-degrading gene expression, nitrite production, and the release of proinflammatory mediators. These protective effects were generally more pronounced with the higher treatment dose. Analysis of publicly available human chondrocyte RNA-sequencing datasets provided complementary support for the relevance of several investigated inflammatory and catabolic targets. Overall, these findings demonstrate that BA protects primary human chondrocytes against inflammatory and catabolic injury and support its further investigation as a potential chondroprotective approach in osteoarthritis.

## Introduction

Articular cartilage is a specialized connective tissue that covers the articulating surfaces of bones within synovial joints [1]. It is avascular and contains chondrocytes, which, under physiological conditions, synthesize extracellular matrix (ECM) components to preserve the structural integrity and biomechanical properties of cartilage [2].

Osteoarthritis (OA) is one of the most prevalent musculoskeletal disorders, particularly among older adults, and represents a major contributor to disability, reduced quality of life, and socioeconomic burden [3]. Clinically, OA is characterized by joint pain, stiffness, and impaired function or mobility [4]. Pathologically, it is a multifactorial, whole-joint disorder characterized by progressive articular cartilage degradation, abnormal subchondral bone remodeling, and synovial inflammation [5]. Effective treatment remains challenging partly because the avascularity and low cellularity of cartilage limit its intrinsic repair capacity, while its dense ECM restricts the penetration of therapeutic agents [6].

Current nonsurgical OA management primarily targets symptom relief and functional improvement through therapeutic exercise, weight management, and pharmacological treatment, particularly topical or oral nonsteroidal anti-inflammatory drugs (NSAIDs); selective cyclooxygenase-2 (COX-2) inhibitors may be considered based on individual gastrointestinal and cardiovascular risk [7, 8]. Although NSAIDs reduce pain and inflammation by inhibiting cyclooxygenase activity and prostaglandin synthesis, prolonged use is associated with gastrointestinal, renal, and cardiovascular adverse effects [7, 8]. Accordingly, inflammation-related pathways, including inducible nitric oxide synthase (iNOS) and nuclear factor kappa B (NF-κB) signaling, are being investigated as potential therapeutic targets [5, 9].

Lipopolysaccharide (LPS), a major endotoxin component of the outer membrane of Gram-negative bacteria, plays a central role in innate immune activation [10]. LPS is recognized by the TLR4–MD-2 receptor complex, triggering MyD88 and TRIF dependent intracellular signaling that activates pro-inflammatory pathways, including NF-κB [11–13]. This process leads to the production of cytokines, including interleukin-1β (IL-1β), interleukin-6 (IL-6), interleukin-8 (IL-8), and tumor necrosis factor-α (TNF-α), along with matrix metalloproteinases (MMPs), which collectively promote inflammation and cartilage degradation [14, 15]. NF-κB activation also induces iNOS expression and consequently increases nitric oxide (NO) production [16]. Excessive NO production has been associated with inflammatory and catabolic responses involved in OA progression [9]. Because of these effects, LPS stimulation is used as a reductionist in vitro model of OA-relevant chondrocyte inflammatory injury [15, 17].

Boron is a naturally occurring trace element found in minerals and compounds such as borax, boric acid (BA), and borates rather than in elemental form [18]. Under physiological conditions, boron is present in humans predominantly as BA; although its essentiality has not been established, nutritional amounts of boron may exert beneficial physiological effects [19, 20]. Available evidence suggests that boron may support bone growth and maintenance and influence vitamin D and steroid hormone metabolism, whereas BA and other boron-containing compounds may promote wound repair [21, 22]. Beyond these metabolic roles, selected boron-containing compounds have shown anti-inflammatory and antioxidant effects, while boron-based strategies are also being investigated in cancer-related and other biomedical applications [19, 20]. Among these compounds, BA has demonstrated anti-inflammatory effects in an experimental rat model of knee OA [23]. However, its effects on OA-relevant inflammatory responses and apoptosis in primary human chondrocytes remain insufficiently understood.

This study aimed to evaluate the anti-inflammatory and anti-apoptotic effects of BA in an in vitro model of OA-relevant inflammatory injury using primary human articular chondrocytes isolated from knee cartilage. Specifically, we investigated whether the most tolerable non-cytotoxic dosage and duration of BA modulates LPS-induced inflammatory, matrix-degrading, and apoptotic responses by assessing selected cytokines, matrix-remodeling mediators, NF-κB pathway-related genes, cellular metabolic activity, cytotoxicity, and apoptosis. To determine whether the selected molecular markers exhibited reproducible regulation in independent human chondrocyte inflammatory models, we additionally performed a targeted cross-study meta-analysis of publicly available RNA-seq datasets. This analysis was designed as external transcriptomic support for the experimental findings rather than as a gene-discovery procedure.

## Material and Method

### Collection of Cartilage Specimen

Articular cartilage specimens were obtained from five patients undergoing total knee arthroplasty (TKA) at Shariati Hospital, Tehran, Iran. Patients with active infections or systemic inflammatory diseases were excluded from the study. Immediately after excision, the specimens were placed in sterile phosphate-buffered saline (PBS) and transported under aseptic conditions to the cell culture laboratory. All participants were informed of the purpose of the study and provided written informed consent before specimen collection. The study protocol was reviewed and approved by the Ethics Committee of Shariati Hospital, Tehran University of Medical Sciences (approval no. IR.TUMS.SHARIATI.REC.1403.067).

### Chondrocyte Isolation

Cartilage specimens were washed three times with sterile PBS and finely minced with sterile scalpels into fragments measuring approximately 1 mm in diameter. The tissue fragments were then digested for 12-16 h in sterile collagenase type II solution prepared in Dulbecco’s modified Eagle’s medium (DMEM) (220 U/mL; Gibco, Thermo Fisher Scientific, catalog no. 17101015, Waltham, MA, USA) at 37 °C under 5% CO_2_ with gentle agitation at 200 rpm. Following digestion, the cell suspension was filtered through a sterile 40-µm cell strainer to remove undigested tissue. The filtrate was centrifuged at 300 × g for 5 min. The resulting cell pellet was washed three times with PBS by resuspension followed by centrifugation at 300 × g for 5 min after each wash. The number of viable cells in each sample was determined using the trypan blue exclusion assay (BIO-IDEA, catalog no. BI-1803-01, Tehran, Iran) using a hemocytometer.

### Chondrocyte Culture

Isolated chondrocytes were seeded in tissue culture-treated 25 cm^2^ culture flasks at an initial density of 3,000 cells/cm^2^. Cells were maintained in high-glucose DMEM (catalog no. 11965175, Gibco, Thermo Fisher Scientific, Waltham, MA, USA) supplemented with 10% (v/v) fetal bovine serum (FBS; catalog no. A5670201, Gibco, Thermo Fisher Scientific, Waltham, MA, USA), and 1% (v/v) penicillin– streptomycin (catalog no. BI-1203, BIO-IDEA, Tehran, Iran). Cultures were incubated at 37 °C in a humidified atmosphere containing 5% CO_2_. Half of the culture medium was replaced every 2 days. Following expansion, chondrocytes at passages 1–3 were used in this study.

### BA Concentration Selection

Primary human chondrocytes were treated with different concentrations of BA (Sigma-Aldrich, St. Louis, MO, USA; catalog no. B6768) at concentrations of 10, 25, 50, 100, 200, 400, and 800 µM and 1.6, 3.2, 6.4, 12.8, and 25.6 mM for 12, 24, 36, or 48 h. Cellular metabolic activity was evaluated using the 3-(4,5-dimethylthiazol-2-yl)-2,5-diphenyltetrazolium bromide (MTT) assay as described in the cell viability assay subsection. Based on this preliminary dose and time response assessment, 100 and 200 µM BA and a 24 h exposure period were selected for the subsequent experiments.

### Experimental Groups and Treatment Protocol

Inflammatory stimulation was induced using LPS (Sigma-Aldrich, St. Louis, MO, USA; catalog no. 437620). Primary human chondrocytes were assigned to four experimental groups: (1) untreated control; (2) cells stimulated with 100 ng/mL LPS for 23 h; (3) cells pretreated with 100 µM BA for 1 h and subsequently stimulated with 100 ng/mL LPS for 23 h without medium replacement; and (4) cells pretreated with 200 µM BA for 1 h and subsequently stimulated with 100 ng/mL LPS for 23 h without medium replacement.

### Cell Viability Assay

Chondrocyte viability was assessed using the MTT assay according to the manufacturer’s instructions, with modifications (DNAbiotech, catalog no. DMA300, Tehran, Iran). Cells were seeded at a density of 7,500 cells per well in tissue culture-treated, clear, flat-bottom 96-well plates and cultured in high-glucose DMEM (catalog no. 11965175, Gibco, Thermo Fisher Scientific, Waltham, MA, USA) supplemented with 20% (v/v) FBS (catalog no. A5670201, Gibco, Thermo Fisher Scientific, Waltham, MA, USA) at a final volume of 100 µL per well. Cultures were maintained at 37 °C in a humidified atmosphere containing 5% CO_2_.

The wells were assigned to four experimental groups. The culture medium was removed, and the cells were washed with PBS before treatment as described in the Experimental Groups and Treatment Protocol subsection. Following treatment, the cells were washed twice with PBS, and 100 µL of MTT working solution (0.5 mg/mL) was added to each well. The plates were incubated for 4 h at 37 °C. The MTT solution was then carefully removed, and formazan crystals were solubilized by adding 100 µL of detergent reagent supplied with the kit to each well, followed by gentle shaking for 10 min at room temperature. For each donor, each experimental condition was evaluated in three replicate wells.

Absorbance was measured at 570 nm, using 690 nm as the reference wavelength, with a PlateLCDread microplate reader (BGT BioGenTechnologies GmbH, Steinfurt, Germany). Relative cell viability was calculated as follows:

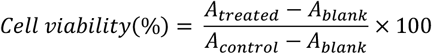

where *A*_*treated*_, *A*_*control*_, and *A*_*blank*_ represent the absorbance values of treated, control, and cell-free blank wells, respectively.

### Lactate Dehydrogenase Assay

Cytotoxicity and loss of chondrocyte plasma membrane integrity were evaluated by quantifying lactate dehydrogenase (LDH) activity released into the culture medium. Chondrocytes from each group were seeded at 1 × 10^4^ cells/well in tissue culture-treated 96-well plates and allowed to adhere for 12 h. The cells were then treated as described in the Experimental Groups and Treatment Protocol subsection. LDH activity was measured using a Cytotoxicity Detection Kit (LDH; Roche Diagnostics GmbH, catalog no. 11644793001, Mannheim, Germany) according to the manufacturer’s instructions.

A background control (200 µL assay medium), a low control (100 µL cell suspension plus 100 µL assay medium), and a high control (100 µL cell suspension plus 100 µL of 2% Triton X-100 solution) were included in each experiment. Briefly, 100 µL of cell-free culture supernatant from each sample was transferred to the corresponding well of an optically clear, flat-bottom 96-well plate. An equal volume (100 µL) of freshly prepared reaction mixture was added to each well, and the plate was incubated for 30 min at room temperature while protected from light. Absorbance was measured at 492 nm using a PlateLCDread microplate reader (BGT BioGenTechnologies GmbH, Steinfurt, Germany). After subtracting the background absorbance, percentage cytotoxicity was calculated as follows:

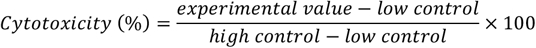

### Apoptosis Assay

Chondrocyte apoptosis was assessed using the Annexin V-FITC/PI Apoptosis Detection Kit (Yas Gene Kowsar, Esfahan, Iran) according to the manufacturer’s instructions. Following treatment, chondrocytes were detached with EDTA-free trypsin and collected by centrifugation at 300 × g for 5 min. The cells were washed twice with cold PBS and resuspended in 1× binding buffer at 1 × 10^6^ cells/100 µL. Cells were stained with 1 µL Annexin V-FITC for 15 min at 25 °C in the dark. Subsequently, 1 µL of propidium iodide (PI) and 400 µL of 1× binding buffer were added. Samples were analyzed using a PAS flow cytometer (Partec GmbH, Münster, Germany). Annexin V-positive/PI-negative and Annexin V-positive/PI-positive cells were classified as early apoptotic and late apoptotic/necrotic cells, respectively.

### Nitric Oxide Production

NO production was estimated by measuring nitrite accumulation in the culture supernatants using a colorimetric NO assay kit (catalog no. KPG-NO, Karmania Pars Gene, Rafsanjan, Iran) according to the manufacturer’s instructions. Briefly, 30 µL of each supernatant was mixed with 100 µL of a freshly prepared mixture of kit solutions A and B and incubated for 10 min at room temperature in the dark. After adding 150 µL of solution E, 150 µL of each reaction mixture was transferred to a 96-well plate. Absorbance was measured at 450 nm using a PlateLCDread microplate reader (BGT BioGenTechnologies GmbH, Steinfurt, Germany). Nitrite concentrations were calculated using the kit-provided standard curve (6.25–100 µM).

### Candidate Gene Selection

Osteoarthritis-associated proteins were identified using STRING version 12.0 by querying the disease term “osteoarthritis” and restricting the organism to *Homo sapiens* (accessed December 14, 2025) [24]. A full STRING functional-association network was generated using a medium-confidence interaction threshold (combined score ≥ 0.400), with text-mining evidence retained. Edges were displayed according to their supporting evidence sources. Functional enrichment analysis was performed directly in STRING using Gene Ontology Biological Process, KEGG and WikiPathways annotations. Terms with a Benjamini-Hochberg-adjusted false-discovery rate (FDR) < 0.05 were considered significant. Candidate genes were prioritized based on their contribution to significant enrichment terms and literature-supported relevance to OA-associated inflammation, NF-κB signaling and extracellular-matrix catabolism. The final gene panel was established before experimental assessment and independent validation using public RNA-sequencing datasets.

### Gene Expression Analysis

Total RNA was extracted from chondrocytes using the PsPure Total RNA Extraction Kit (Pishgaman Sanjesh, Tehran, Iran) according to the manufacturer’s instructions. RNA concentration and purity were assessed using a NanoDrop™ 2000/2000c spectrophotometer (Thermo Fisher Scientific, Waltham, MA, USA). First-strand cDNA was synthesized from 1 µg of total RNA using the cDNA Synthesis Kit (catalog no. YT4500, Yekta Tajhiz Azma, Tehran, Iran) according to the manufacturer’s instructions.

Quantitative real-time PCR (qPCR) was performed for the genes listed in Table 1 using a CFX96™ Real-Time PCR Detection System (Bio-Rad Laboratories, Hercules, CA, USA). Each 20 µL reaction contained HOT FIREPol^®^ EvaGreen^®^ qPCR Mix Plus (catalog no. 08-24-00001, Solis BioDyne, Tartu, Estonia) and specific primers, synthesized by Metabion International AG (Planegg, Germany). The amplification protocol consisted of initial polymerase activation at 95 °C for 15 min, followed by 50 cycles of denaturation at 95 °C for 30 s, annealing at 61 °C for 30 s, and extension at 72 °C for 30 s with a final extension at 72 °C for 5 min.

**Table 1.**
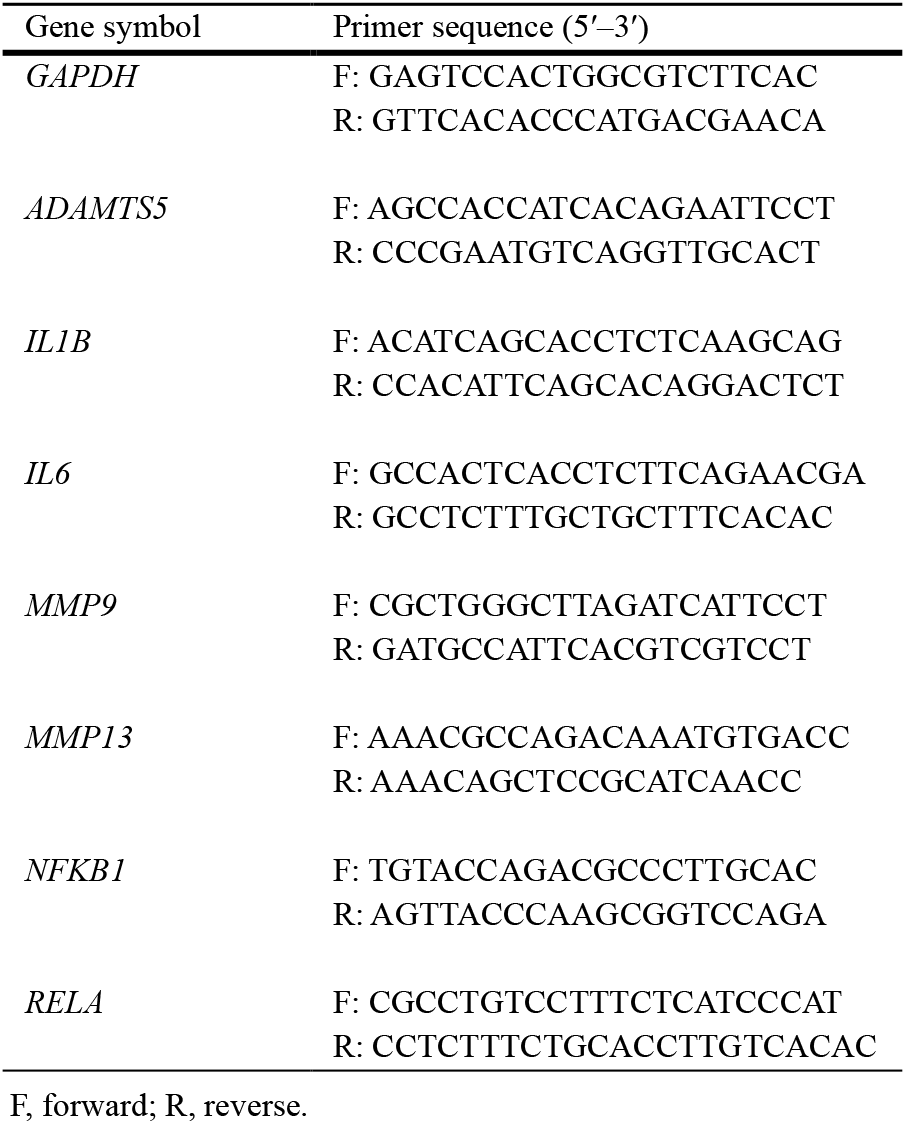
Primer sequences used for quantitative real-time PCR.

### Enzyme-linked Immunosorbent Assay (ELISA)

After the treatment period, culture supernatants were collected and analyzed according to the respective manufacturers’ instructions. Concentrations of IL-6, IL-8/CXCL8, and TNF-α were measured using the Human IL-6 Quantikine QuicKit ELISA (catalog no. QK206), Human IL-8/CXCL8 Quantikine ELISA Kit (catalog no. D8000C), and Human TNF-α Quantikine QuicKit ELISA (catalog no. QK210), respectively (R&D Systems, Minneapolis, MN, USA). MMP-13 concentrations were determined using the Human MMP-13 ELISA Kit (catalog no. EHMMP13; Invitrogen, Thermo Fisher Scientific, Waltham, MA, USA). Optical density was measured at the wavelength specified for each assay using a PlateLCDread microplate reader (BGT BioGenTechnologies GmbH, Steinfurt, Germany), and analyte concentrations were calculated from the corresponding standard curves.

### Public RNA-Seq Meta-Analysis

Publicly available transcriptomic datasets were identified through a targeted search of the NCBI Gene Expression Omnibus (GEO), conducted on October 14, 2025, using combinations of the terms “chondrocyte,” “osteoarthritis,” “IL-1β,” and “RNA sequencing.” Datasets were eligible when they examined primary human chondrocytes, included untreated and IL-1β-treated conditions, provided raw gene-level count data, and contained sufficient biological replication for differential expression analysis. Four datasets met these criteria and were included in the targeted meta-analysis. For GSE215039, only the comparison between IL-1β-treated and untreated OA chondrocytes was included.

Publicly available human chondrocyte RNA-seq datasets were used as an external transcriptomic validation layer for the selected qPCR gene panel. Four GEO datasets with raw count data and relevant inflammatory/OA-relevant stimulation contrasts were included: GSE162510, GSE74220, GSE291878, and GSE215039 [25–28]. For GSE162510, GSE74220, and GSE291878, IL1β-treated samples were compared with untreated or vehicle-treated controls. For GSE215039, IL1β-treated OA chondrocytes were compared with untreated OA chondrocytes.

Statistical analyses were performed in R version 4.5.1 [29]. Differential expression analysis was conducted using DESeq2 version 1.48.1 [30], and random-effects meta-analysis was performed using the metafor package version 5.0-1 [31]. Raw count matrices were analyzed in R using DESeq2. For paired datasets, donor or patient identity was included in the design formula to account for matched samples. Lowly expressed genes were filtered before differential expression analysis. DESeq2 size-factor normalization, dispersion estimation, and negative binomial modeling were then performed. For each dataset, unshrunken log2 fold change estimates and corresponding standard errors were extracted for the predefined target genes *MMP13, MMP9, ADAMTS5, IL1B, IL6, RELA*, and *NFKB1. GAPDH* was excluded because it was used only as the qPCR reference gene.

For each target gene, effect estimates were pooled using an inverse-variance random-effects meta-analysis. The DESeq2 log2 fold change was used as the effect size, and the squared standard error was used as the sampling variance. Between-study variance was estimated using restricted maximum likelihood, and Hartung-Knapp adjustment was applied for confidence interval estimation [32]. The resulting pooled estimates were used to assess whether the selected target genes showed concordant dysregulation across independent inflammatory/OA-relevant chondrocyte transcriptomic datasets.

### Statistical Analysis

Data are presented as mean ± standard deviation (SD) from five biological donors. Technical triplicates were averaged before analysis. Treatment effects were assessed using donor-blocked repeated-measures ANOVA. qPCR data were analyzed on the ΔCt scale, whereas ELISA and Griess data were log_2_-transformed; Greenhouse–Geisser correction was applied where appropriate. The planned within-donor comparisons were evaluated using paired t-tests with Holm adjustment. Tukey HSD was used for qPCR all-pair comparisons. Early/late apoptosis was additionally analyzed using two-factor repeated-measures ANOVA (stage × treatment). Analyses and figures were performed in R (version 4.5.1). P<0.05 was considered statistically significant.

## Results

### BA Protects Chondrocytes Against LPS-Induced Cell Injury

The initial dose selection experiment showed that chondrocyte viability declined as the BA concentration and exposure time increased. Based on these findings, 100 and 200 µM BA were selected for the subsequent experiments (Fig. 1A). LPS significantly reduced cell viability compared with the control group. Treatment with 200 µM BA significantly improved viability, whereas the improvement observed with 100 µM BA did not reach statistical significance (Fig. 1B).

**Fig. 1.**
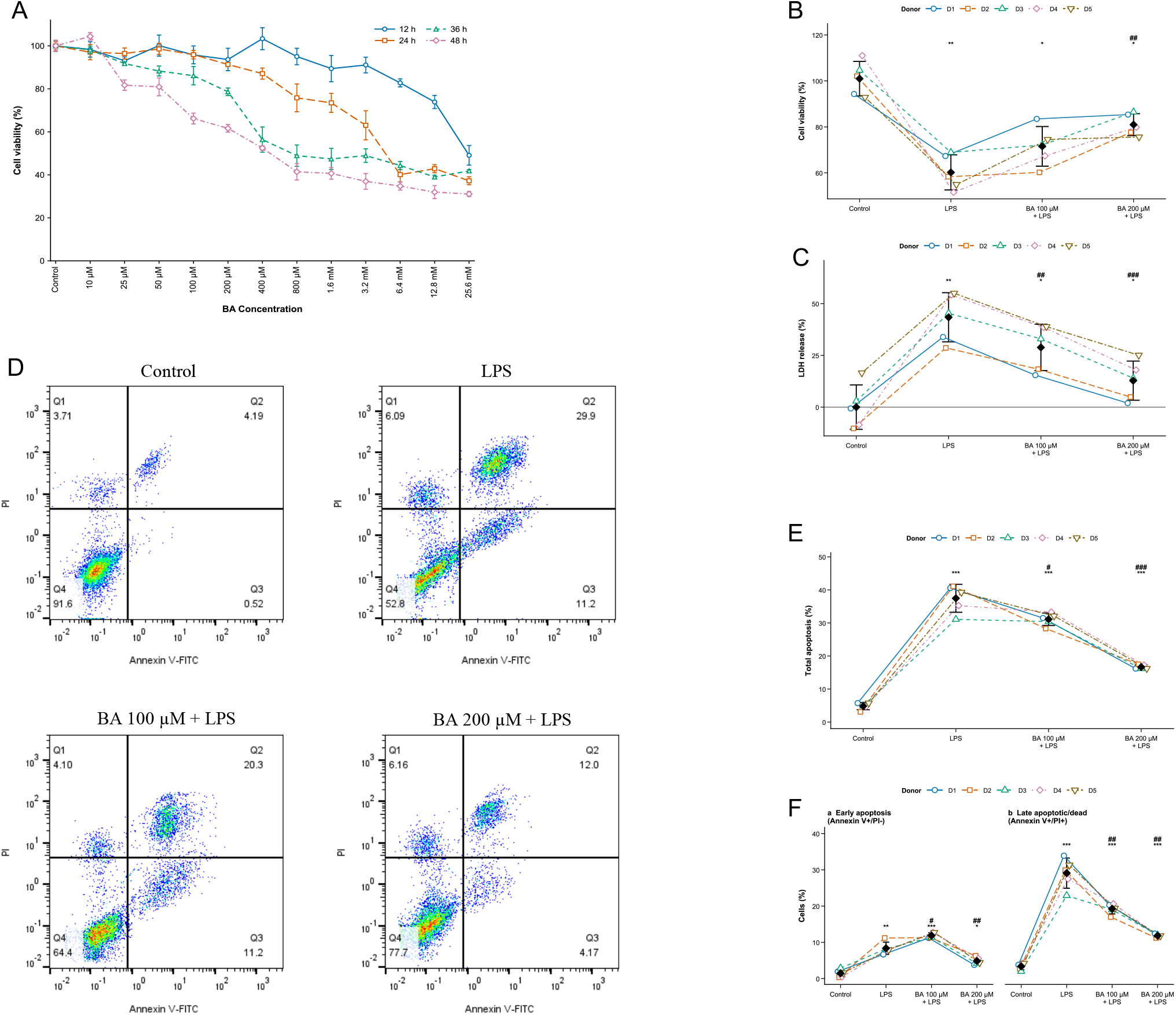
Boric acid (BA) concentration selection and effects on lipopolysaccharide (LPS)-induced loss of viability and apoptosis in human chondrocytes. (**A**) MTT cell viability following exposure to BA. (**B**) MTT cell viability following treatment with LPS (100 ng/mL) alone or with BA for 23 h. (**C**) LDH release assay. (**D**) Representative Annexin V–FITC/ PI flow cytometry plots. (**E**) Total apoptosis. (**F**) Early apoptotic (Annexin V+/PI−) and late apoptotic/dead (Annexin V+/PI+) cells. In (**A**), data are mean ± SD of three technical replicates. In (**B**), (**C**), (**E**) and (**F**), open symbols and connecting lines represent individual donors, and black diamonds represent mean ± SD (n = 5). *p < 0.05, **p < 0.01, and ***p < 0.001 versus control; #p < 0.05, ##p < 0.01, and ###p < 0.001 versus LPS

Flow-cytometric analysis confirmed the protective effect of BA against LPS-induced apoptosis. LPS markedly increased total apoptosis, while both BA concentrations significantly reduced this response (Fig. 1C-D). Examination of the individual apoptotic populations revealed different responses at the two BA concentrations. Treatment with 100 µM BA increased the early apoptotic population but reduced the late apoptotic/dead population compared with LPS alone. In contrast, 200 µM BA significantly reduced both early and late apoptotic/dead populations (Fig. 1E).

LPS also increased LDH release, indicating loss of membrane integrity. Both BA concentrations significantly reduced LDH release, with a greater reduction observed at 200 µM (Fig. 1F). Collectively, these findings demonstrate that BA protects chondrocytes against LPS-induced loss of viability, apoptosis, and membrane damage.

### BA Suppresses LPS-Induced Inflammatory Gene Expression and Nitrite Production

LPS significantly increased the expression of *ADAMTS5, MMP13, MMP9, IL1B*, and *IL6* compared with the control group. Both concentrations of BA significantly attenuated the LPS-induced expression of these genes (Fig. 2A). The inhibitory effect was generally more pronounced at 200 µM. Compared with 100 µM BA, the higher concentration produced further reductions in *ADAMTS5, MMP9*, and *IL1B* expression, whereas the differences in *MMP13* and *IL6* expression were not significant. Neither *NFKB1* nor *RELA* expression differed significantly among the treatment groups.

**Fig. 2.**
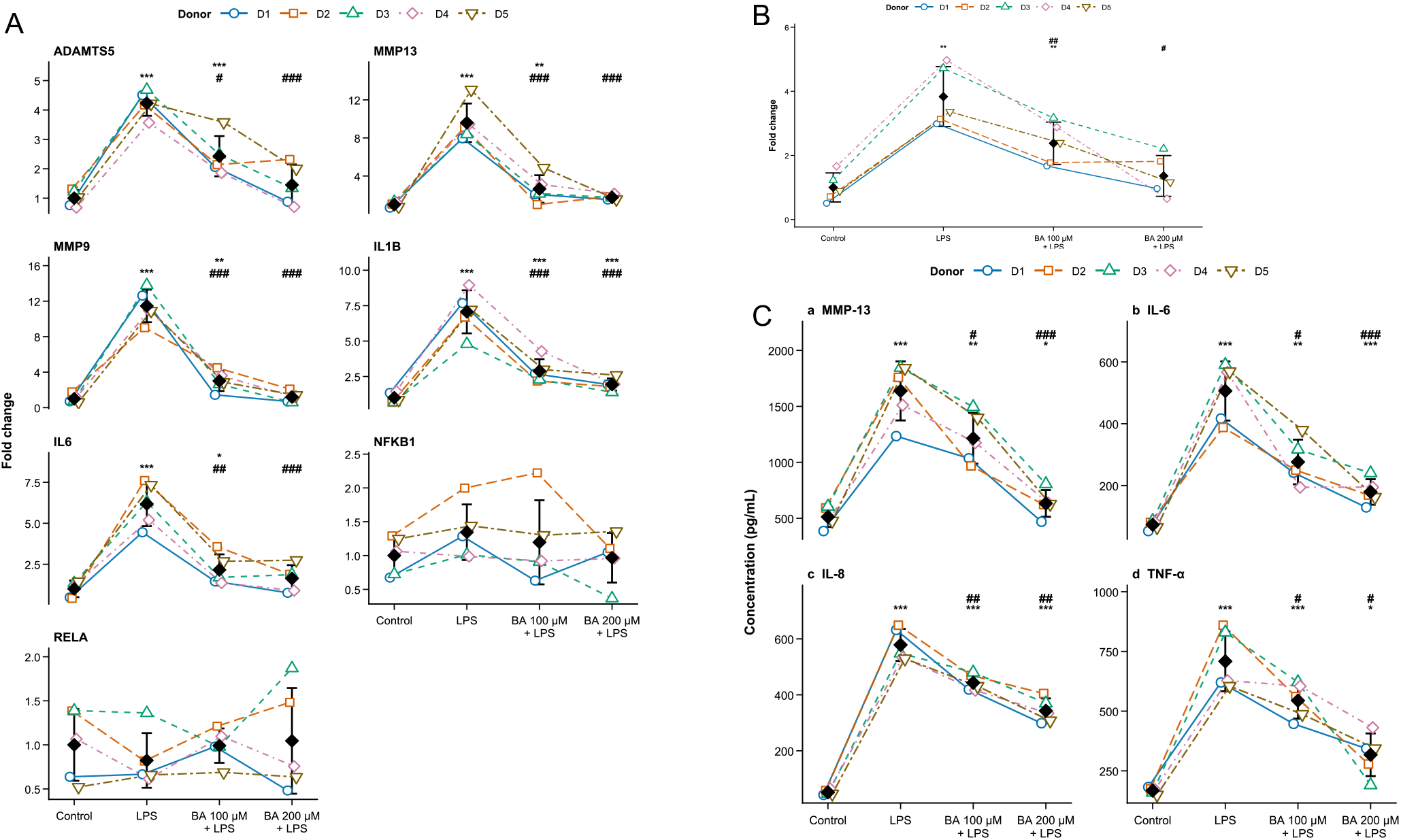
Effects of boric acid (BA) on lipopolysaccharide (LPS)-induced inflammatory responses in human chondrocytes treated with LPS (100 ng/mL) alone or with BA. (**A**) Relative expression of *ADAMTS5, MMP13, MMP9, IL1B, IL6, NFKB1*, and *RELA* measured by quantitative PCR. (**B**) Nitrite production measured by the Griess assay. (**C**) MMP-13, IL-6, IL-8, and TNF-α concentrations in conditioned medium measured by ELISA. Open symbols and connecting lines represent individual donors, and black diamonds represent mean ± SD (n = 5). *p < 0.05, **p < 0.01, and ***p < 0.001 versus control; #p < 0.05, ##p < 0.01, and ###p < 0.001 versus LPS

The Griess assay showed a similar response. LPS markedly increased nitrite production, whereas treatment with BA significantly suppressed this increase. The reduction was more pronounced with 200 µM BA, which brought nitrite production closer to the control level (Fig. 2B). These findings indicate that BA attenuates the inflammatory response induced by LPS.

### BA Reduces the Release of MMP-13 and Pro-inflammatory Cytokines

LPS significantly increased the secretion of MMP-13, IL-6, IL-8, and TNF-α into the conditioned medium. Treatment with BA reduced the release of these inflammatory mediators, with the strongest overall response observed at 200 µM (Fig. 2C). These changes were consistent with the gene expression and nitrite production findings, supporting the anti-inflammatory effect of BA in LPS-stimulated chondrocytes.

### Independent Transcriptomic Support for the Selected Gene Panel

To assess the reproducibility of the selected molecular markers, four independent human chondrocyte RNA-seq datasets were analyzed using random-effects meta-analysis. Positive pooled effects were observed for *MMP13, IL6, IL1B, RELA*, and *NFKB1*, with 95% confidence intervals excluding zero. *MMP9* showed a positive but non-significant pooled effect, whereas *ADAMTS5* did not demonstrate consistent regulation across datasets. Considerable between-study heterogeneity was observed for most genes, indicating variation in effect magnitude among experimental systems despite broad directional concordance for several inflammatory and catabolic markers (Fig. 3; Table 2).

**Table 2.** Characteristics of the publicly available RNA-sequencing datasets included in the targeted meta-analysis.

| GEO accession | Biological source | IL-1β exposure | Contrast included | Samples analyzed | Study design | Sequencing platform |
| --- | --- | --- | --- | --- | --- | --- |
| GSE162510 | Primary human OA chondrocytes | 0.2 ng/mL, 24 h | IL-1β vs untreated control | 10 + 10 | Paired by donor | Illumina NextSeq 500 |
| GSE74220 | Primary human chondrocytes from hip OA patients | 1 ng/mL, 4 h | IL-1β vs untreated control | 3 + 3 | Paired by donor | Illumina HiSeq 2000 |
| GSE291878 | Primary human chondrocytes | 1 ng/mL, 6 h | IL-1β plus vehicle vs vehicle control | 5 + 5 | Paired by donor | Illumina NextSeq 2000 |
| GSE215039 | Primary human knee OA chondrocytes | 1 ng/mL, 24 h | OA IL-1β vs untreated OA | 5 + 5 | Paired by donor | Illumina NovaSeq 6000 |

**Fig. 3.**
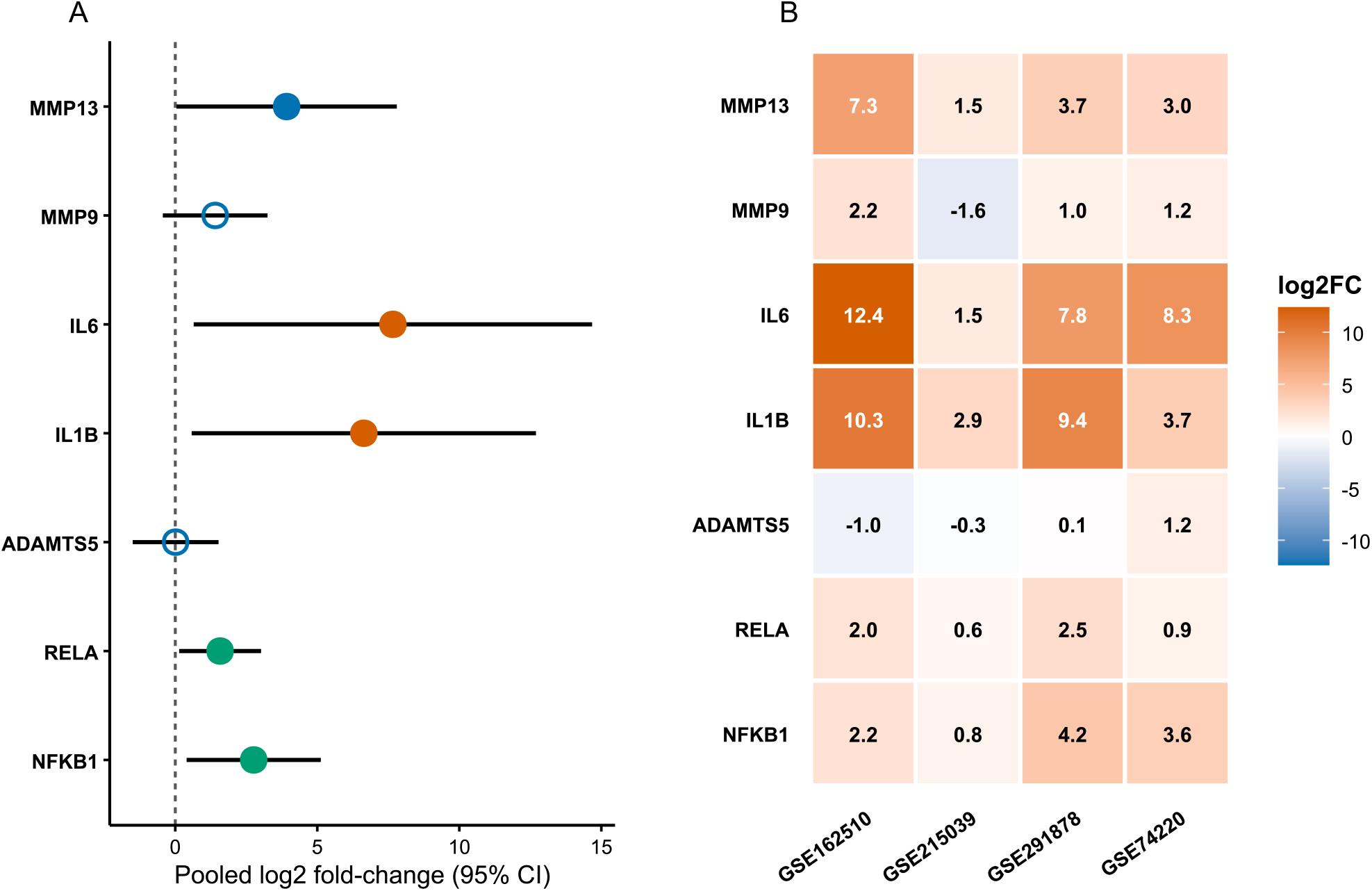
Meta-analysis of inflammatory and catabolic gene expression in IL-1β-stimulated human chondrocytes. (**A**) Random-effects pooled log2 fold-change (log2FC) estimates and 95% confidence intervals for the prespecified seven-gene panel across four independent RNA-sequencing datasets. Models were fitted using inverse-variance weighting, restricted maximum-likelihood estimation of between-study variance, and Hartung-Knapp confidence intervals. Filled symbols indicate confidence intervals that exclude zero, whereas open symbols indicate confidence intervals that include zero. Colors group genes involved in extracellular matrix catabolism, inflammatory cytokine signaling, and NF-κB regulation. (**B**) Heatmap showing study-specific DESeq2 log2FC estimates. Positive and negative values indicate increased and decreased expression, respectively, following IL-1β treatment. GSE215039 represents osteoarthritis (OA) chondrocytes treated with IL-1β versus untreated OA chondrocytes; the remaining datasets compare IL-1β-treated with control chondrocytes.

## Discussion

Osteoarthritis (OA) is a multifactorial whole-joint disorder in which inflammatory activation of articular chondrocytes contributes to progressive cartilage degradation [5, 7]. Pro-inflammatory stimuli promote the production of cytokines, nitric oxide, matrix metalloproteinases, and aggrecanases, thereby disrupting extracellular matrix homeostasis and contributing to chondrocyte dysfunction and death [9, 14, 15]. Lipopolysaccharide (LPS) activates TLR4-dependent signaling and reproduces important inflammatory and catabolic responses in chondrocytes; consequently, it is widely used as an in vitro model of OA-relevant inflammatory injury [11, 17]. Boron-containing compounds, including boric acid (BA), have demonstrated anti-inflammatory, antioxidant, and tissue-protective activities, while BA has also shown beneficial effects in experimental models of OA [22, 23]. BA is particularly relevant because inorganic boron is predominantly present as BA under physiological conditions and is principally absorbed in this form [19, 20]. Nevertheless, whether BA can protect primary human articular chondrocytes against LPS-induced cytotoxicity, apoptosis, inflammation, and matrix-degrading responses remains insufficiently understood [17, 23, 33].

Our initial functional findings indicate that BA attenuated LPS-induced injury in primary human chondrocytes. Preliminary MTT screening showed that cellular metabolic activity declined as BA concentration and exposure duration increased. On this basis, 100 and 200 µM BA and a total BA exposure of 24 h, comprising 1 h of pretreatment followed by 23 h of LPS coincubation, were selected for the main experiments (Fig. 1A). Treatment with LPS alone for 23 h reduced cellular metabolic activity and increased LDH release, indicating impaired cellular function and loss of plasma membrane integrity (Fig. 1B–C). Flow cytometric analysis supported this pattern, as LPS markedly increased total apoptosis, predominantly through an increase in the late apoptotic/dead population (Fig. 1D–F). In the BA groups, chondrocytes were pretreated with 100 or 200 µM BA for 1 h and subsequently coincubated with BA and LPS for 23 h. Compared with LPS alone, both BA concentrations reduced LDH release, apoptosis rate, and the proportion of late apoptotic/dead cells, whereas a significant improvement in MTT metabolic activity was observed only with 200 µM BA. The distribution of apoptotic populations nevertheless revealed a concentration related difference: 100 µM BA increased the proportion of early apoptotic cells relative to LPS alone while reducing late apoptosis/death, whereas 200 µM BA significantly reduced both populations. This pattern may indicate that the lower BA concentration limited advanced membrane compromised injury without fully preventing the initiation of apoptosis; however, measurements at a single time point cannot establish the temporal progression or reversibility of these cellular states. A previous study similarly demonstrated that LPS induces inflammatory, catabolic, and apoptotic injury through TLR4-associated signaling in human chondrocytes [17]. BA has also been reported to suppress LPS-induced TNF-α production in THP-1 monocytes through a thiol-dependent mechanism and to reduce inflammatory and cartilage-catabolic markers in a rat model of knee OA [23, 33]. Collectively, our findings extend this evidence to primary human articular chondrocytes and indicate that BA, particularly at 200 µM, produces a broader cytoprotective response involving cellular metabolic activity, membrane integrity, and apoptosis.

The cytoprotective effects of BA were accompanied by coordinated suppression of inflammatory and matrix-degrading responses in LPS-stimulated chondrocytes. LPS treatment alone strongly increased the expression of *ADAMTS5, MMP13, MMP9, IL1B*, and *IL6*, together with nitrite production and the secretion of MMP-13, IL-6, IL-8, and TNF-α (Fig. 2A–C). These responses are consistent with the capacity of LPS to activate inflammatory and catabolic pathways in human chondrocytes [34]. BA pretreatment followed by LPS coincubation significantly attenuated the LPS-induced expression of all five genes at both BA concentrations, with generally greater inhibition at 200 µM. Compared with 100 µM BA, 200 µM BA produced further reductions in *ADAMTS5, MMP9, MMP13* and *IL6*, whereas the differences between the two BA concentrations were not significant for *IL1B*. BA also reduced nitrite accumulation and the secretion of MMP-13, IL-6, IL-8, and TNF-α, with the strongest overall response observed at 200 µM. The concordant reductions in *MMP13* and *IL6* transcripts and their corresponding secreted proteins, together with the consistent responses across qPCR, Griess, and ELISA measurements, support a coordinated anti-inflammatory and anti-catabolic effect rather than the alteration of a single marker. MMP-13 and ADAMTS-5 are major mediators of collagen and aggrecan degradation, respectively, while pro-inflammatory cytokines and excessive nitric oxide reinforce catabolic signaling, impair matrix synthesis, and contribute to chondrocyte dysfunction in OA [9, 14, 35]. Neither *NFKB1* nor *RELA*, which encode components of the canonical NF-κB complex, differed significantly among the experimental groups. Because NF-κB activity is principally controlled through events such as IκBα phosphorylation and degradation and the subsequent nuclear translocation of existing NF-κB dimers, unchanged *NFKB1* and *RELA* transcript levels do not exclude pathway activation [12]. Nevertheless, NF-κB or MAPK signaling cannot be identified as the mechanism of BA action because IκBα phosphorylation, NF-κB nuclear translocation, and MAPK activation were not directly assessed. Previous studies similarly showed that BA suppressed LPS-induced TNF-α production in THP-1 monocytes and reduced inflammatory and cartilage-catabolic markers in experimental knee OA [23, 33]. Collectively, these findings indicate that BA limits multiple components of the inflammatory and catabolic chondrocyte response; however, because extracellular matrix content or synthesis was not measured, reduced degradative-mediator expression should not be interpreted as direct evidence of preserved or restored cartilage matrix.

To determine whether the selected molecular response panel was relevant beyond our experimental system, we integrated four independent human chondrocyte RNA-seq datasets using random-effects meta-analysis. Positive pooled effects with 95% confidence intervals excluding zero were observed for *MMP13, IL6, IL1B, RELA*, and *NFKB1* (Fig. 3; Table 2). The pooled regulation of *MMP13, IL6*, and *IL1B* was concordant with their induction by LPS in our primary chondrocytes, supporting the relevance of these inflammatory and catabolic markers across independent human chondrocyte models. In contrast, *MMP9* showed a positive but nonsignificant pooled effect, whereas *ADAMTS5* was not consistently regulated across the included datasets. Considerable between-study heterogeneity was observed for most genes, likely reflecting differences in donor characteristics, OA status, stimulation conditions, exposure duration, and experimental design. The positive pooled effects for *RELA* and *NFKB1* also differed from their unchanged expression in our LPS-stimulated cells, further indicating that transcriptional regulation of NF-κB components is context dependent and should not be equated with pathway activation [12]. Importantly, the public datasets examined IL-1β-stimulated chondrocytes and excluded BA treatment; therefore, this analysis does not independently validate the effects of BA or establish its mechanism of action. Rather, it provides complementary evidence that several components of our predefined gene panel are broadly responsive in inflammatory and OA-relevant human chondrocyte systems, while also showing that *MMP9* and *ADAMTS5* require more context specific interpretation. Together with the experimental findings, the meta-analysis strengthens the biological rationale for the selected response panel, whereas conclusions regarding BA remain grounded in the primary LPS experiments.

Viewed together, the findings indicate that BA influenced several connected components of the chondrocyte response rather than a single experimental endpoint. The use of primary cells from five donors under matched treatment conditions, allowed the effects on metabolic activity, membrane integrity, apoptosis, gene expression, nitrite production, and secreted mediators to be evaluated within the same biological framework. The generally stronger response at 200 µM, combined with the decline in cellular metabolic activity at higher concentrations and longer exposures during the preliminary screening, suggests that the protective effects of BA occur within a defined exposure range. This concentration sensitivity is consistent with the context dependent effects reported in cartilage and joint models [23, 36, 37]. In a rat model of knee OA, BA reduced IL-1β and TNF-α levels and MMP-13 activity/expression and improved cartilage histopathology, while intra-articular BA improved macroscopic and histological repair scores in a rabbit osteochondral-defect model [23, 36]. Conversely, an earlier study in embryonic chick cartilage found that BA decreased proteoglycan and collagen synthesis while increasing macromolecule release and endoprotease activity [37]. Differences in species, developmental stage, concentration, metabolic conditions, and the presence or absence of inflammatory stimulation may account for these divergent responses [37,36,23]. Therefore, the present findings extend the available evidence to primary human articular chondrocytes, but they do not establish direct preservation of cartilage matrix or predict a therapeutic effect in the human joint.

This study has several limitations. Although the repeated-measures design accounted for within donor variability, cartilage was obtained from five patients at a single center, which limits the precision and generalizability of the findings. Expansion of primary chondrocytes in monolayer culture can alter their differentiated phenotype, and LPS stimulation cannot reproduce the interactions among cartilage, synovium, subchondral bone, immune cells, and mechanical loading that characterize OA in vivo [38,7]. In addition, BA was administered as a 1 h pretreatment followed by 23 h of coincubation with LPS; consequently, the study primarily evaluates protection against an impending inflammatory challenge rather than reversal of an established response. Only two BA concentrations, one LPS concentration, and one principal assessment time were examined in the main experiments, and BA-only groups were not included in the principal LPS comparisons. Direct measurements of extracellular matrix synthesis or loss were not performed, and the study did not assess NF-κB or MAPK activation, intracellular oxidative stress, mitochondrial function, or apoptosis-related signaling proteins. The public RNA-seq datasets involved IL-1β stimulation and did not contain BA treated samples, so they cannot independently confirm the effects of BA. Finally, concentrations effective in cell culture cannot be directly extrapolated to achievable systemic or intra-articular exposures in humans.

Overall, BA pretreatment followed by coincubation with LPS attenuated multiple features of LPS-induced injury in primary human articular chondrocytes, including loss of cellular metabolic activity, membrane damage, apoptosis, inflammatory mediator production, and expression of matrix-degrading factors. The response was more consistent at 200 µM, although the preliminary concentration screening also showed that higher or more prolonged exposure was not uniformly beneficial. These findings support cytoprotective, anti-inflammatory, and anti-catabolic activity under the conditions examined, but they do not identify the underlying signaling mechanism or demonstrate cartilage preservation. Further studies should test BA after inflammatory activation has been established, examine pathway activity and redox regulation directly, and determine whether the observed molecular effects are maintained in 3D cultures, human cartilage explants, and multicellular joint models. Pharmacokinetic studies will also be needed to establish whether biologically active concentrations can be achieved safely in vivo. Within these boundaries, the present study provides a basis for continued investigation of BA as a potential chondroprotective compound in OA.

## Conclusions

This study demonstrated that BA attenuated LPS-induced injury in primary human articular chondrocytes. BA reduced the loss of cellular metabolic activity, membrane damage, apoptosis, nitrite production, inflammatory mediator secretion, and expression of matrix-degrading factors. The RNA-seq meta-analysis supported the biological relevance of several investigated markers across independent inflammatory human chondrocyte models but did not independently validate the effects of BA. These findings identify BA as a potential cytoprotective, anti-inflammatory, and anti-catabolic compound under the experimental conditions examined. Further studies should clarify its mechanism of action, evaluate treatment after inflammatory injury has been established, and determine whether these effects translate into matrix preservation in cartilage explants, 3D models, and in vivo systems.

## Acknowledgments

This study was fully funded by the Center for Orthopedic Trans-Disciplinary Applied Research.

## Data Availability

The data presented in this study will not be publicly available prior to the publication of the research findings but may be obtained from the corresponding author upon reasonable request.

